# Territory establishment, song learning strategies and survival in song sparrows

**DOI:** 10.1101/804021

**Authors:** Çağlar Akçay, S. Elizabeth. Campbell, Saethra Darling, Michael D. Beecher

## Abstract

In most songbirds the processes of song learning and territory establishment overlap in the early life of young birds who usually winds up with songs matching those of their territorial neighbors in their first breeding season. In the present study, we examined the relationships among the timing of territory establishment, the pattern of song learning and territorial success in a sedentary population of song sparrows (*Melospiza melodia*). Males in this population show high song sharing within neighborhoods derving from their learning most of their songs from neighboring males. These shared songs are preferentially used in interactions with neighbors. Males also show significant variation in the timing of territory establishment, ranging from their first summer until the next spring. Using a three-year dataset, we found that the timing of territory establishment did not systematically affect the composition of the song repertoire of the tutee: early establishers and late establishers learned equally as much from their primary tutors, and had a similar number of tutors and repertoire size. Timing of territory establishment also did not have an effect on subsequent survival on territory. Therefore, the song learning program of song sparrows seems versatile enough to lead to high song sharing even for birds that establish territories relatively late.

## 1. INTRODUCTION

Bird song is unusual among vocal communication systems, because it is one of the few in which animals learn their vocal signals. Vocal learning has been found only in humans, songbirds (oscine passerines), cetaceans, bats and two other orders of birds (parrots and hummingbirds) (Baptista & Schuchmann, 1990; Boughman, 1998; Pepperberg, 1994; Reiss & McCowan, 1993; Todt, 1975). Aside from human language, songbirds have provided the main model system for studying social learning of communicative signals (Brainard & Doupe, 2002; Doupe & Kuhl, 1999).

In most songbirds, particularly in close-ended learners, the young bird learns its songs after it has dispersed from its natal area and is prospecting for and establishing a breeding territory in a new area (Beecher, Campbell, & Stoddard, 1994; DeWolfe, Baptista, & Petrinovich, 1989). Indeed, in most species of songbirds where song learning has been studied in the field, the learning process results in the young bird having learned their songs from their eventual neighbors (Beecher et al., 1994; Liu & Kroodsma, 2006; Nelson & Poesel, 2009; Payne, 1983; Wheelwright et al., 2008).

If the function of a song learning strategy is to learn songs of your eventual neighbors, the timing of learning should evolve such that the bird memorizes songs and actively shapes its song repertoire while establishing a territory amongst its tutor-neighbors. But early laboratory studies found that song memorization is usually limited to a short period early in the natal summer (Marler & Peters, 1977, 1987, 1988). This poses a dilemma for species that are close-ended learners (birds that learn their repertoire of songs in their first year of life and subsequently don’t modify it): the young bird will need to learn the songs of its eventual neighbors, but will have to learn these songs in advance of knowing exactly who these neighbors and what their songs will be.

A partial solution to this dilemma was proposed by Nelson and Marler (1994). According to their selective attrition model, a young bird memorizes songs only in its natal summer. The following spring, however, when the young birds are singing and trying to establish their territories, they produce more songs than they will eventually keep. During this time of overproduction, they interact with older, territorial birds, and prune their song repertoire down to just those songs that best match those of these older birds who will be their neighbors. Field and laboratory studies have provided some support for the over-production and selective attrition model in several species (Nelson, 1992; Nelson & Marler, 1994; Nelson, Marler, & Morton, 1996; Nelson & Poesel, 2009; Nordby, Campbell, & Beecher, 2007).

Another solution to the dilemma is flexibility in the timing of the memorization phase depending on the young bird’s social experience. Evidence for such flexibility comes from studies on marsh wrens (*Cistothorus palustris*) (Kroodsma & Pickert, 1980, 1984). Young male marsh wrens who hatch early in the summer and are exposed to recorded marsh wren songs in the laboratory in the summer and following spring will complete their song learning in the first 60-80 days of life (Kroodsma, 1978). In a laboratory study using recorded song, Kroodsma and Pickert (1980) compared the tendency of marsh wrens to memorize new songs in the spring as a function of whether they had been raised on a photoperiod simulating early hatching (June) or late hatching (August). Both groups received the same regimes of song tutoring in the natal year, but when exposed to new songs the following spring, only the late (August) birds added some of these new songs to their repertoires. This suggests that late hatching and under-exposure to song may extend the period in which a bird is capable of memorizing new songs into the following spring.

In this paper we examine the relationship of song learning and the timing of territory establishment in a resident population of song sparrows (*Melospiza melodia*). Song sparrows are a temperate songbird species in which only males sing; female song is very rare (Arcese, Stoddard, & Hiebert, 1988; Beecher, Campbell et al. unpublished observations). Male song sparrows are close-ended learners that develop a repertoire of about 9 songs (range: 5-13 songs) in their first year of life, which they do not modify in subsequent years (Nordby, Campbell, & Beecher, 2002).

Several field studies (Nordby et al 1999; Akcay et al 2014) have identified key features of song learning in our study population of song sparrows. (1) Each young bird (hereafter tutee) learns from several older males (hereafter tutors). (2) The final repertoire of the tutee is biased towards a single, primary tutor who on average accounts for about half of the tutee’s song repertoire, although this proportion varies from 0.3 to 1. This primary tutor is usually an immediate neighbor (Nordby, Campbell, & Beecher, 1999). (3) Song learning begins in the natal summer, but ultimately the bird learns (or retains) more songs from tutors who survive into the next spring than from tutors who don’t. (4) Song sharing with neighbors is important in territorial interactions in this population as the birds use shared songs in a hierarchical, graded signaling system (Akçay, Tom, Campbell, & Beecher, 2013; Beecher, Stoddard, Campbell, & Horning, 1996; Burt, Campbell, & Beecher, 2001; Stoddard, Beecher, Campbell, & Horning, 1992). (5) Although it is not completely clear how song sharing benefits a bird, the degree of song sharing with neighbors in the young bird’s first breeding season is positively correlated with the number of years the bird survives on territory (Beecher, Campbell, & Nordby, 2000; P. L. Wilson, Towner, & Vehrencamp, 2000). In contrast, repertoire size is unrelated to a bird’s territory tenure (Beecher et al., 2000).

Early laboratory studies of song learning in male song sparrows in an eastern population (*M. melodia melodia* subspecies) revealed a sensitive period mostly limited to the first summer (Marler & Peters, 1987). Subsequent laboratory studies of our resident population (*morphna* subspecies) have found that learning of new songs can occur in the fall and following spring as well (Nordby, Campbell, & Beecher, 2001; Nulty et al., 2010; Templeton, Burt, et al., 2012) although how common this ‘late learning’ is in nature is still unknown, and generalizing to natural conditions from laboratory studies is difficult).

In our field site and nearby resident populations, young song sparrows are observed to ‘float’ over a wide area until they manage to find a territory (Arcese, 1987, 1989; J. N. M. Smith & Arcese, 1989; Templeton, Reed, Campbell, & Beecher, 2012), which can happen from as early as July (at about 2-3 months of age) to as late as April of the following spring. As suggested above, this individual variation in territory establishment might have consequences for song learning, especially given that song sparrows in our population sing throughout the late summer and fall (though at a reduced rate) which means that potentially tutees can overhear singing interactions among territorial adult males as well as directly interact with them.

An early establishing male, particularly those establishing territories in the summer (July and August) would likely have more opportunities earlier to interact with their primary tutors, leading potentially to more song learning to them. However such an advantage would only materialize if the primary tutor also survived into the breeding season, given our finding that birds learn (or retain) more songs from tutors that survive the winter (past January 1) than from tutors who do not (Akçay, Campbell, Reed, & Beecher, 2014; Nordby et al., 1999). Depending on the year, about 30 to 40% of the older birds do not survive over winter. Thus, we might expect that the effect of early territory establishment would only be seen in cases where the primary tutor has also survived into the breeding season.

In contrast, a late establisher may be at a disadvantage in terms of learning the neighbors’ songs. Our earlier studies suggest that in at least some cases, young birds establish their territories late because they have been shut out of the area where they learned songs in their natal summer, as can happen, for example, when none of the tutor-neighbors die over winter (Nordby et al., 1999). In this case the songs the young bird memorized in his natal year will generally be poor matches to the songs of their new neighborhood, leaving the young bird with the alternatives of learning a new set of songs in short order or of just retaining, through selective attrition, the best-matching of his early-memorized songs. Other late establishers may simply have hatched late and will not have heard enough song in their natal summer, in which case they would have to do much of their song learning in the fall or following spring, which might lead to sub-optimal repertoires. A late establisher may have fewer opportunities to engage in direct interactions or overhear interactions between neighboring males, and these are thought to have an important role in the attrition process (Nelson & Marler, 1994). Another potential handicap for late establishers is that even young song sparrows singing non-crystallized song are treated more aggressively by territorial adults in the spring than they are in the fall, and more aggressively in the fall than they are in the summer (Templeton, Campbell, & Beecher, 2012). Moreover, while young song sparrows are seen associating closely with older males in the summer, they are not seen doing so in the spring (Templeton, Reed, et al., 2012).

In the present study we analyze a three-year dataset on the timing of territory establishment by young birds in our resident population of song sparrows. In three consecutive years starting with 2009, we banded and recorded young males during the period of their song learning and attempted to track their time of territory establishment through systematic surveys. We then compared their song repertoires to those of all adult males in the population in the bird’s first year of life, and attempted to correlate the characteristics of song learning in birds establishing their territories in the natal summer, fall or following spring. We also compared the tutees subsequent survival on territory to test whether early establishing birds were either somehow of better quality and therefore survived for longer on territory. This pattern may be expected if timing in territory establishment is correlated with competitive ability of the young bird as has been proposed for migratory birds (Kokko, 1999). If that is the case earlier establishing birds should survive more years on territory than later establishing birds. Alternatively, establishing your territory early be provide an advantage in itself, which would also lead to early establishing birds surving longer on territory.

## 2. METHODS

### 2.1. Study site and subjects

This study is part of a long-term study of song sparrows located in Discovery Park, Seattle, Washington, USA, that started in 1986 (Beecher et al., 1994). More information on the specifics of the site can be found in Beecher (2008). This population is resident year-round and males generally defend their territories all year, with the exception of during molting (August) and cold weather periods in November-December, when birds show reduced territoriality (but often are still on their territory). Breeding usually starts in March or April depending on the weather conditions, particularly the El Nino cycle (S. Wilson & Arcese, 2003), although song sparrows start becoming territorial again and singing immediately after the winter solstice when days start to lengthen (G. T. Smith, Brenowitz, Beecher, & Wingfield, 1997). We therefore considered January 1 as the starting point of Spring. Each year between 120 and 150 adult males hold territories in the portion of the park under study. Males were caught with mist nets or Potter traps and each male was fitted with a US Fish and Wildlife Service metal band and three color bands for visual identification. Often multiple juveniles were caught in the same net by herding them into the net as a flock (Templeton, Reed, et al., 2012).

### 2.2 Surveying

During the study, we kept track of the arrival and disappearance of males on territory by visiting territories every two weeks throughout the year, with the exceptions noted above of August (molting) and November and December (inclement weather) when we surveyed the study area opportunistically and banded new birds whenever we could. We used either playbacks or observation of singing males to determine whether a territory holder was still present or had been replaced by a new bird.

We counted a bird as territorial if he was observed singing on a territory and approached playback of conspecific song. We took the date of territory establishment of the young bird to be the date of first such observation on a territory if the same area was known to have been held by another bird recently (within a few weeks) and the young bird kept the territory into the spring. In our study site, all suitable areas for song sparrow territories are occupied at any given point in time, and we have rarely observed song sparrows expanding into “no-man’s land” areas where there were no song sparrows previously, except for cases where habitat had been significantly changed (e.g. planting or growth of new shrubbery). We classified territory establishment dates into three categories: the summer of the hatch year (before September), fall of hatch year (September to January) and next spring (January to April, the majority of these birds established their territories in January and February).

In total we determined territory establishment dates and song learning for 71 young birds who hatched in the years 2009 (n=30), 2010 (n=22) and 2011 (n=19) and established territories in our study site sometime between the summer of their hatch year and the subsequent spring. The majority of the subjects (44 out of 71, 61.9%) were banded either in the summer with juvenile plumage or in the fall with breeding plumage but singing plastic song. The rest of the subjects were banded after January 1^st^ of their second calendar year with breeding plumage but were identified as second year birds from their songs which still showed plastic elements (see Figure 1 for examples of song development at different stages in the first year).

**Figure 1.**
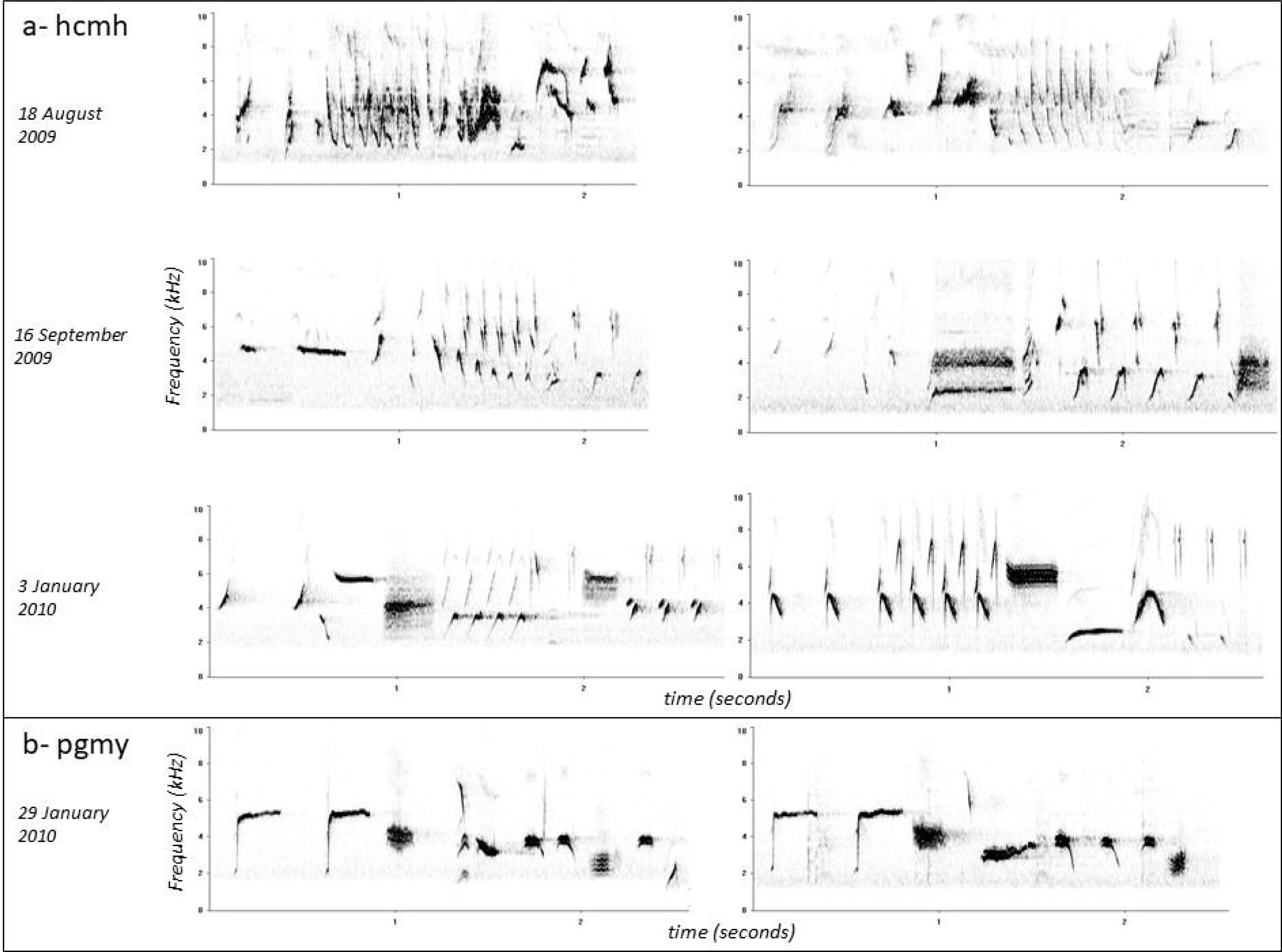
Examples of song development in summer (top row), fall (middle row) and spring (bottom two rows). The songs in panel a were recorded from a single male (hcmh) who was observed to be territorial starting in late July, a few weeks after being banded in juvenile plumage. The two example songs in each row were recorded in the same recording session on the indicated date and are meant to give the reader a sense of the range in development of the songs. Note that by fall this bird had a rather well developed repertoire in which song types could be distinguished although the songs still plastic as can be seen by the non-consistent repetition of intro notes and the trill notes. By January, the bird sang fairly crystallized songs where notes were repeated with high consistency and the overall song structure did not vary from one rendition to the next (although occasionally he would sing plastic song). The two songs in panel b were recorded from a different male (pgmy), also banded in the preceding July, but not seen to be territorial until January. Note that the songs of this male are more plastic at this stage than hcmh as indicated by the non-consistent repetition of the elements within and across song renditions.

We tracked the survival of birds through 2016 by surveying the area as indicated above. Song sparrows don’t move significant distances once they establish a territory, moving at most a few territories over (Hughes & Hyman, 2011). We therefore considered birds to have lost territory if they were not observed on their prior territory, any of the nearby territories or anywhere else in our study area, and if their original territory was being defended by another male (Akçay, Campbell, & Beecher, 2015).

### 2.3 Song recording and song learning analyses

We recorded the repertoire of young males and their potential tutors with a digital recorder and a shotgun microphone (Marantz PMD 660 and Sennheiser ME66/K6). We considered the repertoire fully recorded after at least 16 switches in a continuous recording, which has been shown to be a large enough sample to capture the entire repertoire (Nordby et al., 2002). From these recordings we carried out song analyses as in our previous studies (Akçay et al., 2014; Beecher et al., 1994; Nordby et al., 1999). Briefly, we made spectrograms of each song in the repertoire of each tutee and potential tutor using Syrinx (John Burt, www.syrinxpc.com). We considered each adult male that held a territory in June of the tutee’s hatch year as a potential tutor. Three judges independently compared the visual match between the songs of the tutee and potential tutors. After this stage, the three judges compared their matches and arrived at a consensus sheet where all judges agreed upon the matches.

If a male was implicated as having the sole best match to a tutee song, he was given a score of 1 (full credit) for that song. For songs where more than one male was judged to have the best match, the score was split among these males (e.g. if there were two males, each received 0.5, etc.). Split-credit songs like these happen because of high levels of song sharing within neighborhoods in our population (Hill, Campbell, Nordby, Burt, & Beecher, 1999). For about half of the songs, tutorship was shared in this way (46.5% in 2009 cohort; Akçay et al., 2014). For each tutee, the tutor with the highest tutoring score was defined as the primary tutor. For this primary tutor, we noted whether he survived past January 1^st^ of the second calendar year of the tutee.

### 2.4. Data analyses

From the dates of territory establishment, we classified each tutee as having established a territory in the natal summer, natal fall, or spring (after January 1^st^ of their second year). Our dependent variables were repertoire size, proportion of songs learned from the primary tutor and number of tutors that accounted for the entire repertoire. We analyzed these dependent variables with linear mixed models with territory establishment season, tutor survival into spring and their interaction as the predictor variables and cohort as a random factor. We analyzed survival of the tutees with a general linear mixed model with Poisson distribution and log-link, adding cohort as a random factor and territory establishment season as a fixed factor. The analyses were carried out in R (R Core Team, 2012)..

## 3. RESULTS

Fourteen (15.6%) and 32 (35.6%) of the tutees were first observed to be territorial in the natal summer and fall respectively, while 44 tutees (48.9%) were first observed to be territorial in spring after January 1^st^(11 of these established their territories in March and April while the rest established their territories in January and February). The season in which the tutees established their territory did not have a significant effect on either the proportion of songs learned from the primary tutor, the number of tutors, or the overall repertoire size (Table 1, Figure 2). Whether the primary tutor survived into the first spring of the tutee had a significant effect on proportion of the repertoire this tutor accounted for: tutees whose primary tutors survived past January 1^st^ learned a higher proportion of their repertoire from them than did tutees whose primary tutor did not, and had a smaller number of tutors, replicating earlier findings in our population. There also was no interaction effect of territory establishment season and whether the primary tutor survived into spring. Finally, there was no difference in survival depending on the timing of first territory establishment (*χ*^2^=1.70, p=0.43, Figure 3).

**Table 1.** The results of linear mixed models with tutee territory establishment season, primary tutor’s survival into the first spring of the tutee and their interaction as fixed factors and cohort as a random factor. The reported values are *χ*^2^ values and the associated p-values from Wald tests.

|  | Proportion learned from primary tutor | Repertoire size | Number of tutors |
| --- | --- | --- | --- |
| Season of territory establishment | 0.08 (0.96) | 1.22 (0.54) | 1.86 (0.39) |
| Primary tutor's survival into January | 4.70 (0.026) | 0.31 (0.58) | 1.77 (0.18) |
| Interaction | 0.34 (0.84) | 0.04 (0.97) | 1.19 (0.55) |

**Figure 2.**
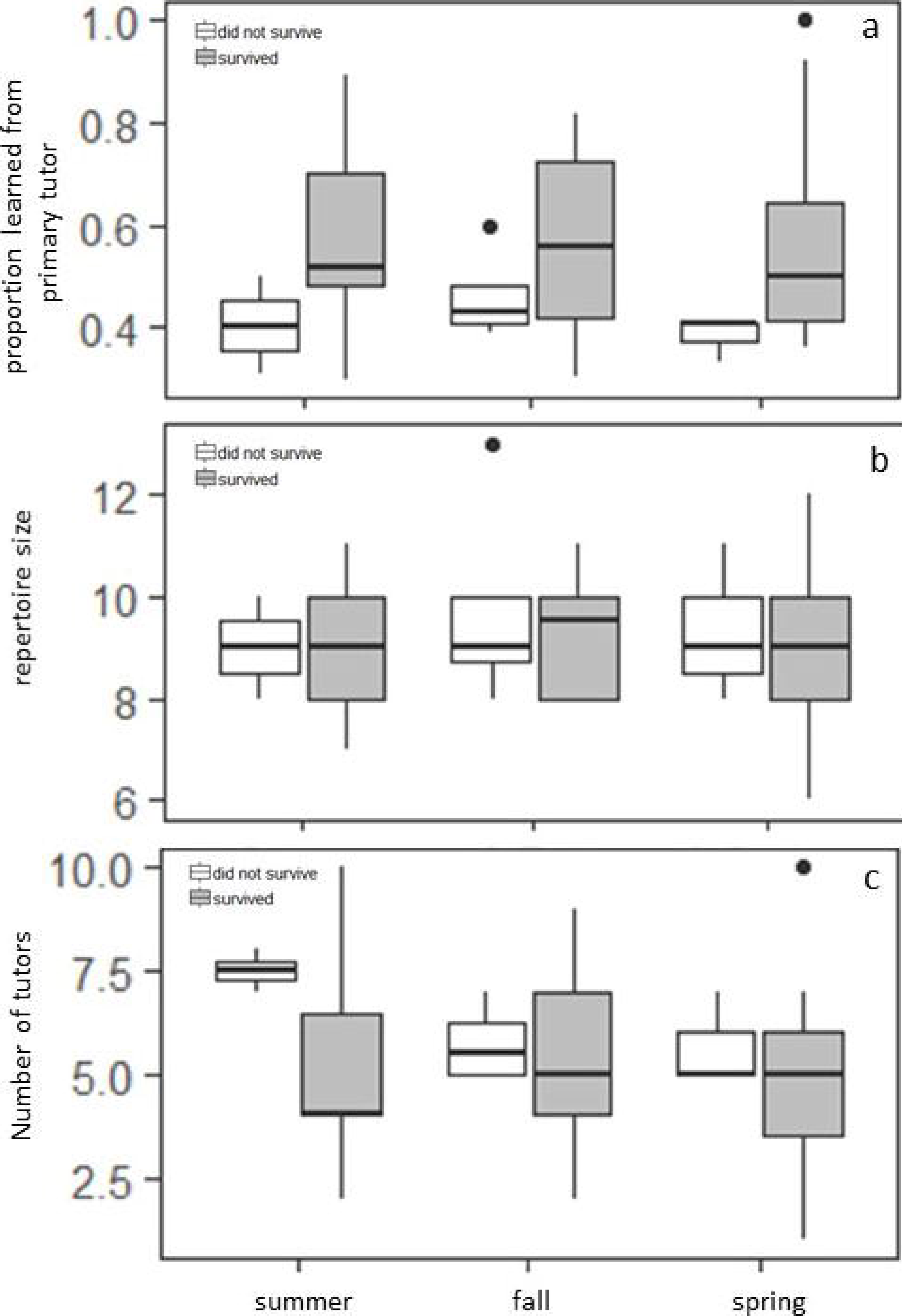
Proportion of songs learned from the primary tutor (a), repertoire size (b) and number of tutors tutees had (c) depending on territory establishment date and whether the primary tutor survived into the first spring of the tutee (white= tutor did not survive, gray= tutor survived).

**Figure 3.**
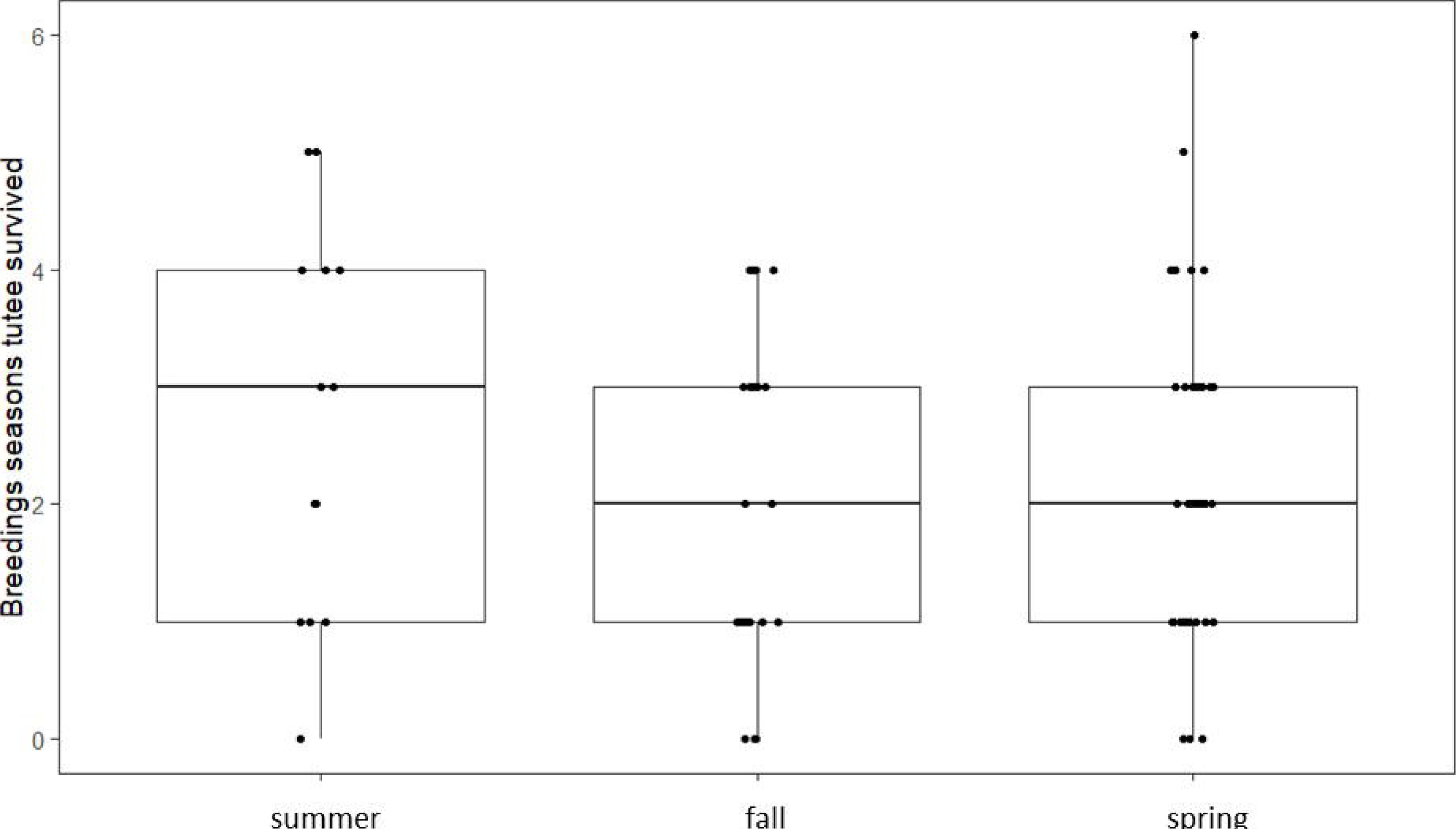
Number of years survived on territory depending on the timing of territory establishment.

## 4. DISCUSSION

We tested whether the timing of territory establishment has any influence on song learning strategies of male song sparrows. While we replicated our earlier finding that tutor survival into spring increases the tutoring influence of that bird, we found that whether the young bird established his territory in the summer, fall or spring did not affect either proportion of the song repertoire learned from that tutor, the number of tutors or the final repertoire size of the tutee. We also found no support that early establishing birds experienced a fitness benefit in the long-term as survival on territory did not significantly differ between birds establishing their territories in different seasons.

### 4.1 Timing of Song Learning and Territory Establishment

The fact that timing of territory establishment does not affect these aspects of song learning suggests at least two possibilities. First, at least some birds that establish their territories late may have been present all along. It is hard to detect juvenile song sparrows that are not territorial as they float around and therefore while not territorial, the birds may have been present and listening in on the singing interactions between adults. Given that eavesdropping in such a way is also a potent factor in song learning (Beecher, Burt, O'Loghlen, Templeton, & Campbell, 2007; Templeton, Akçay, Campbell, & Beecher, 2010), this may compensate for the lack of direct interactions with adults that they would have if they were territorial. Relatedly, young song sparrows may be able to memorize a great number of songs from many males during their floater period, such that even late-establishing birds are still able to produce the songs of an older bird that becomes a primary tutor (instead of trying to reach a repertoire that is composed of a single or few songs from many tutors). Radio-tracking studies of young birds in our study population show that they sometimes cover a large amount of ground (Templeton, Reed, et al., 2012), and during these movements they are likely to overhear many adults singing. Given laboratory studies that failed to detect an upper limit in recognition memory for songs in song sparrows (Stoddard, Beecher, Loesche, & Campbell, 1992), it is plausible that a bird is able to memorize a very large number of songs during this floater phase. This hypothesis predicts that late establishers would be initially singing larger repertoires which they would then winnow down to a repertoire that matches their primary tutor and other neighbors. Note that this strategy of memorization and production of a large repertoire may lead to a cost in terms of the quality of learning. For instance in Puget-Sound white-crowned sparrows (*Zonotrichia leucophrys pugetensis*), young birds that sang large repertoires (e.g. 4 songs) imitated the local songs more poorly than young birds that sang the species-typical repertoire size of a single song (Nelson & Poesel, 2014). Although we did not quantify the match between the tutee and tutor version of the songs, it is possible that late establishers may match their primary tutor songs less well compared to early establishers.

A second, non-exclusive possibility is that song sparrows can memorize new songs until at least the time they establish their territories, whenever this happens. Estimates of the timing of memorization phase comes from laboratory studies of hand-raised birds, typically tutored with recorded songs played from speakers—the so-called “tape tutors” (Marler & Peters, 1987, 1988). Under more naturalistic conditions involving either live birds or song presentation that represent naturalistic bouts of singing, these estimates have proven to significantly underestimate the period of song memorization which often extends into the first spring of the birds (Baptista & Petrinovich, 1984; Kroodsma & Pickert, 1980, 1984; Nordby et al., 2001; Nulty et al., 2010; Templeton, Burt, et al., 2012). Furthermore as discussed above there is evidence that the memorization phase in some species, like marsh wrens and white-crowned sparrows show flexibility with respect to hatching date and the presence of live tutors (Baptista & Petrinovich, 1984, 1986; Kroodsma & Pickert, 1980, 1984; Petrinovich & Baptista, 1987). Presumably, interactions with live tutors in the lab simulate territorial interactions in the field. Therefore, the finding of flexibility in memorization phase suggests that the sensitive period for song memorization would close in the summer for birds establishing at that time and close the following spring for birds establishing at that time.

### 4.2. Song learning in sedentary and migratory populations

Our population of song sparrows are sedentary with birds singing year-round. This may result in the evolution of a memorization phase that is longer. The situation would be different for migratory birds which often travel to a wintering ground where they either don’t hear conspecific song, or hear songs that are not local to the areas to which they will return next spring. Thus, memorization of songs during the first winter may not be adaptive, and birds should ignore or avoid memorizing these songs. Nevertheless, song memorization phase might reopen the next spring. Some evidence on this point comes from the previously-mentioned study of marsh wrens (Kroodsma and Pickert, 1980; 1984) and a study of chaffinches (Thielcke & Krome, 1991). In the chaffinch study, the authors tutored juvenile birds caught in late summer and early Fall with one song in the fall and winter and another song in the next spring. Despite the fact that the birds had significant song exposure in the field before being caught, they nevertheless learned the spring song from tape tutors. None of the birds copied the fall song however, suggesting that chaffinches may be insensitive to song presented in this period. The Nulty et al. study (2010) cited earlier also found evidence in our population that tutors that were heard in summer and fall were less effective tutors compared to tutors heard in summer and spring, suggesting that song sparrows too may be insensitive to fall song.

Although our population is sedentary, some nearby populations of song sparrows display altitudinal migratory strategies in which the birds migrate from their high-altitude, snow covered breeding grounds to lower altitudes where they would hear sedentary song sparrows singing that are not necessarily local to their population. In a study of one such population Hill et al. (1999) found that levels of song sharing was high and not different than the sedentary population that we studied here, suggesting that song learning programs are not likely to be different in this migratory population. Given that final territories in this population are not established until next spring, this suggest that memorization phase either stays open throughout fall and spring or reopens in the spring. To distinguish these possibilities, a common garden experiment comparing the memorization of songs presented in different seasons to birds from these sedentary and migratory populations would be required.

### 4.3. Timing of territory establishment and fitness

We found no evidence that early establishing males experienced a long-term fitness benefit compared to later establishing males. Given that in our population, almost all areas that is suitable for song sparrows is occupied, the variation in timing of territory establishment is likely to be due to opening of vacancies due to the disappearance of a prior territory owner or the presence of a large territory that may be split into two territories enough to support two song sparrow pairs. The latter cases may arise over spring: later in the spring, when breeding is ongoing, birds who are depredated often are not replaced by new arrivals but their territories are taken over by their existing neighbors (Akçay et al., 2012). This fact means by summer time, there are often territories that used to hold two territories but are now defended by a single song sparrow, and juveniles may be able to insert themselves into these territories more easily.

Most studies on timing of territorial behavior and subsequent fitness effects has been on migratory species which arrive at their breeding territories from somewhere else (Brooke, 1979; Francis & Cooke, 1986; Kokko, 1999; Lozano, Perreault, & Lemon, 1996). Variation in timing of territory establishment has been studied in year-round resident species less often (Dixon, 1956; Matthysen, 1989). In one such study, Matthysen (1989) did not detect any differences in fitness (in terms of survival) and eventual territory quality between early- and late-establishing males in the year round resident European nuthatches (*Sitta europea*), consistent with our present findings.

Variation in the timing of territory establishment has been studied in a partially migratory population of eastern song sparrows in rural Pennsylvania (Hughes & Hyman, 2011). In this population, some males established their territories early in the Spring while about ¼ of males established territories later (after the first nest has hatched). Hughes and Hyman (2011) found that timing of first-time territory establishment did not relate to fitness in terms of survival. Indeed, late establishing males who subsequently moved to a different, non-overlapping territory had the highest reproductive success in terms of number of nests and young fledged. These results too suggest that late establishing males do not necessarily consist of low-quality individuals, consistent with the present results.

### 4.4 Conclusions

In conclusion, in a large multi-year dataset we found no effect of timing of territory establishment on song learning strategies as exemplified by the three main parameters that vary across individual song sparrows: proportion of repertoire learned from the primary tutor, repertoire size and how many tutors the tutee learned his songs from. These results indicate that song sparrows are able to match the songs of their neighbors even if they establish their territories late, suggesting a versatile learning program either due to a longer memorization phase, a large number of songs memorized or practicing singing while not being territorial. We also found no evidence that late establishing males suffer a fitness cost, consistent with earlier findings in year-round resident species. Future research can experimentally manipulate song experience of birds in the field in realistic ways (e.g. Mennill et al., 2018) while keeping track of their individual histories to dissociate the different hypotheses regarding how birds can adaptively shape their learned song repertoire and territorial strategies.

## Acknowledgements

We thank Veronica Reed and Chris Templeton for help with field work.

